# Timely excision of prophage Φ13 is essential for the *Staphylococcus aureus* infectious process

**DOI:** 10.1101/2025.02.04.636441

**Authors:** Olivier Poupel, Gérald Kenanian, Lhousseine Touqui, Charlotte Abrial, Tarek Msadek, Sarah Dubrac

**Affiliations:** Institut Pasteur, Université Paris Cité, CNRS UMR2001, Department of Microbiology, Biology of Gram-Positive Pathogens Unit, F-75015 Paris, France; Institut Pasteur, Université Paris Cité, Department of Global Health, Cystic Fibrosis and Bronchial Diseases Unit, F-75015 Paris, France; Institut Pasteur, Université Paris Cité, CNRS UMR6047, Department of Microbiology, Stress Adaptation and Metabolism in Enterobacteria Unit, F-75015 Paris, France

## Abstract

Mobile genetic elements play an essential part in the infectious process of major pathogens, yet the role of prophage dynamics in *Staphylococcus aureus* pathogenesis is still not well understood. Here we studied the impact of the Φ13 *hlb-* converting prophage, whose integration inactivates the *hlb* β-toxin gene, on staphylococcal pathogenesis. We showed that prophage Φ13 is lost in approximately half the bacterial population during the course of infection. Inactivation of the Φ13 *int* recombinase gene, essential for insertion/excision, locked the prophage in the bacterial chromosome, leading to a significant loss of virulence in a murine systemic infection model. In contrast, the non-lysogen strain (ΔΦ13), where the *hlb* beta-hemolysin gene is reconstituted, displayed strongly increased virulence. Accordingly, histopathological analyses revealed more severe nephritis in mice infected with bacteria lacking prophage Φ13 (ΔΦ13), compared to infection with the parental strain. Infection with the Δ*int* mutant, where beta-hemolysin production is abolished, led to the least severe renal lesions. Cytokine induction in a human neutrophil model showed significantly increased IL-6 expression following infection with the beta-hemolysin producing strain (ΔΦ13). Our results indicate that timely *in vivo* excision of the Φ13 prophage is essential for progression of the *Staphylococcus aureus* infectious process: early excision leads to rapid host death whereas the inability to excise the prophage significantly reduces staphylococcal virulence.

**IMPORTANCE:** This study highlights prophage Φ13 excision as a critical factor in *S. aureus* pathogenesis, influencing infection outcomes by balancing rapid host killing with reduced bacterial virulence. This mechanism may represent a bet-hedging strategy in genetic regulation, resulting in a mixed bacterial population capable of rapidly switching between two processes: bacterial colonization and host damage. Unraveling this dynamic opens new possibilities for developing targeted therapies to disrupt or modulate prophage activity, offering a novel approach to mitigating *S. aureus* infections.

## INTRODUCTION

*Staphylococcus aureus*, a major human pathogen, causes severe invasive infectious diseases targeting every type of organ, such as pneumonia, endocarditis, osteomyelitis or septicemia. Yet it is also a commensal, asymptomatically colonizing up to 30% of the human population, with the transition between the two lifestyles poorly understood (1). Mobile genetic elements make up approximately 20% of its genome, leading to variations in gene combinations that can strongly influence *S. aureus* virulence (2). Among these, temperate bacteriophages play an important role in genetic variability. Φ13, a member of the Sa3int Siphoviridae *S. aureus* bacteriophage family, is responsible for a dual phenotypic conversion (3, 4) and is by far the most widespread among human clinical isolates (5). Its chromosomal attachment site (*attB*) lies within the coding sequence of the hemolysin B gene, *hlb*, encoding the neutral sphingomyelinase C β-toxin, such that it acts as a phage regulatory switch (6). Φ13 lysogeny leads to disruption of *hlb* and loss of β-hemolytic activity, with excision reconstituting an intact *hlb* gene (5, 7, 8).

Phage Φ13 also carries the *iec* immune evasion cluster including several genes involved in escape from the host innate immune response: *scn*, *chp* and *sak* (9, 10). Natural mobility has been shown in vivo for Φ13-related phages during murine skin colonization and rabbit invasive infection as well as in isolates from cystic fibrosis patients and healthy individuals (5, 11–13). Additionally, among *S*. *aureus* isolates from recurrent furunculosis, 45% produced β-hemolysin compared to only 19% of those from nasal colonization (14).

Given the importance of phage Φ13 excision on β-hemolytic activity, we wished to study the effect of the presence or absence of phage Φ13 on *S*. *aureus* virulence during systemic infection in three different configurations that had not been previously studied:

- a) preventing phage excision and consequently β-hemolytic activity
- b) absence of phage Φ13 leading to continuous β-hemolytic activity
- c) simultaneous presence of phage Φ13 and a functional *hlb* gene.

We show that timely excision of the Φ13 prophage is essential for pathogenesis, with early β-hemolysin production leading to a significant increase in virulence and elevated levels of the pro-inflammatory cytokine IL-6. Conversely, preventing phage Φ13 excision significantly lowers staphylococcal virulence.

## RESULTS

### Mobility dynamics of *Staphylococcus aureus* prophages

*S. aureus* strain HG001, a derivative of sepsis isolate NCTC 8325 (15), carries prophages Φ11, Φ12 and Φ13 (7, 16). Specific oligonucleotide primer pairs were designed to detect the presence or absence of each prophage (Supplementary Table 2). Using chromosomal DNA from a closely related strain cured of all phages, SH1000, an amplicon corresponding to the native bacterial locus without Φ13 (oligonucleotide pair OMA7/OMA9, hybridizing on either side of the *attB* site, Supplementary Fig. 1A) was detected, but no amplification occurred with oligonucleotides specific for Φ13 insertion (OMA7/OMA8; Supplementary Fig. 1B). With the HG001 Φ13 lysogen strain, both types of amplicon were generated by PCR, suggesting spontaneous Φ13 excision within the bacterial population (Supplementary Fig. 1B). We quantified relative prophage levels within the bacterial population by qRT-PCR during growth under optimal laboratory conditions (TSB at 37°C with aeration). Our results indicate that prophage Φ11 is highly mobile with approximately 10% of cells lacking the phage whereas Φ12 was much more stable with less than 0.01% of non-lysogen cells (Fig. 1A). Interestingly, the Φ13 prophage was lost in about 0.5% of the population under these growth conditions (Fig. 1A).

**Figure 1:**
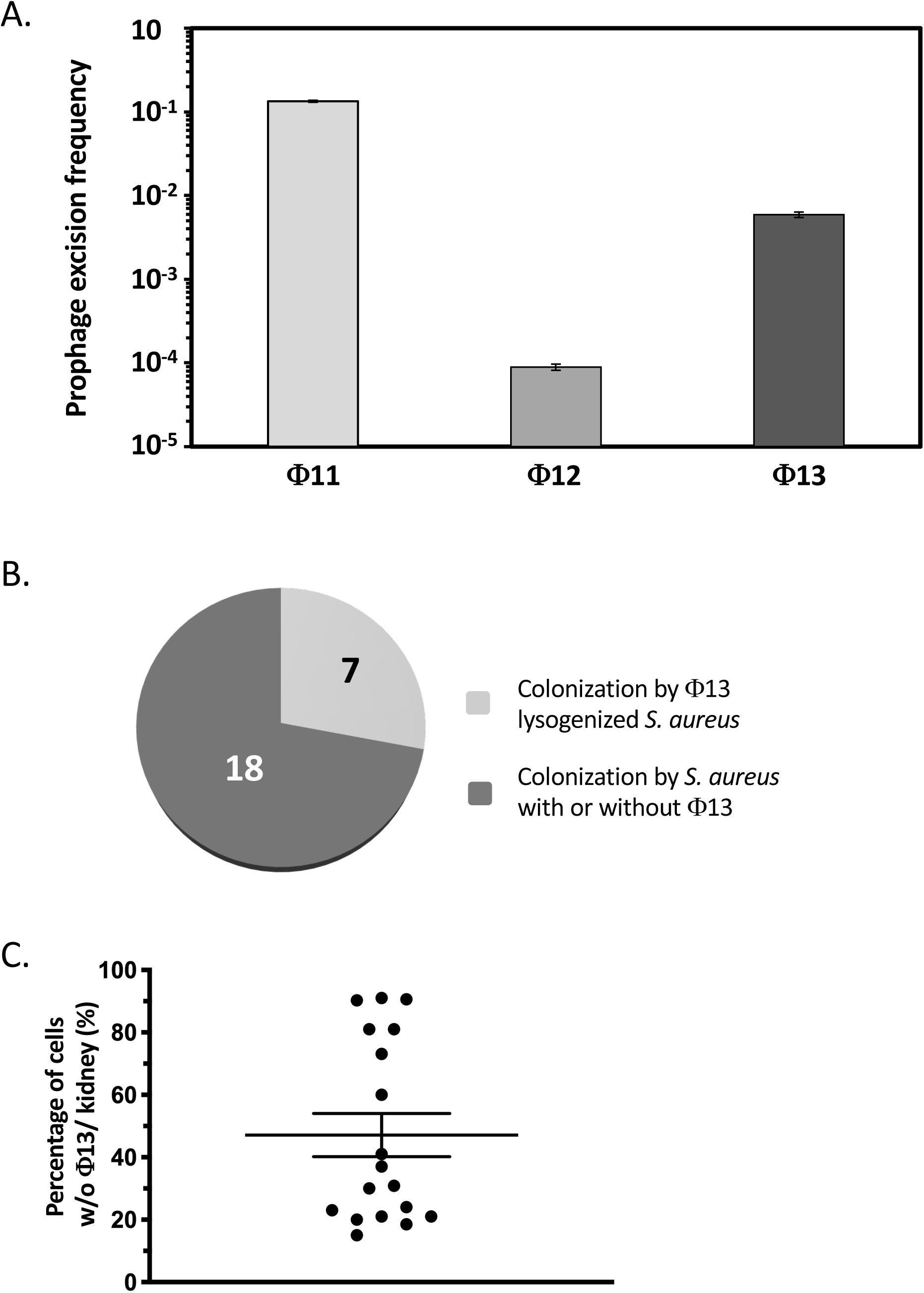
Prophage Φ13 excision in vitro and in vivo. A) Prophage excision frequencies in strain HG001. qRT-PCR quantification of Φ11, 12, and 13 prophage content in *S*. *aureus* strain HG001 grown in TSB at 37°C with aeration. Specific primer pairs are described in Supplementary Table 2. The results are the mean of 4 technical replicates and are representative of 3 independent biological replicates. B) Quantification of Φ13 lysogens in bacterial populations from infected kidneys. Mice were infected from the caudal lateral vein with a sub-lethal dose (2 × 10^7^ c.f.u.) of *S. aureus* strain HG001 Φ13::*aad*(9). Mice were sacrificed four days post-infection and kidneys harvested for c.f.u. counts on TSA and TSA with spectinomycin. 25 infected kidneys were examined and the proportion of homogeneous lysogen population (less than 0.5% non-lysogen) and heterogeneous bacterial population with a mix of lysogen and non-lysogen bacteria is shown. C) Proportion of non-lysogen bacteria per kidney in organs infected with a mixed population of lysogenic and non-lysogenic bacteria.

### Excision of prophage Φ13 is increased *in vivo*

As shown above, prophage Φ13 is spontaneously excised during growth under laboratory conditions. Since this phage carries genes encoding the SAK staphylokinase virulence factor, as well as SCIN and CHIP, involved in escape from the innate immune response (17), we examined Φ13 excision frequency during the infectious process.

We generated a strain carrying the *aad*(9) spectinomycin resistance gene inserted within the Φ13 genome of the HG001 strain (see Materials and Methods, and Supplementary Table 1 strain ST1387). We verified that the *aad*(9) cassette did not interfere with Φ13 stability during growth in vitro. In the ST1387 strain (HG001Φ13 - *aad*(9)), the prophage excision frequency was 0.58% ± 0.05%, which is comparable to the excision rate observed in the wild-type HG001 strain (0.59% ± 0.05%), as determined by qRT-PCR. Four days following sub-lethal (2 × 10^7^ c.f.u.) intravenous inoculation with *S. aureus* strain ST1387, mice were sacrificed and kidneys collected for c.f.u. counts. Out of 25 harvested kidneys, 18 were colonized by a mixed bacterial population with respect to Φ13 lysogeny (Fig.1B). In some mice a single kidney was colonized with a mixed population with the other only by Φ13 lysogens, suggesting that loss of Φ13 is a *post facto* event to kidney colonization. The average loss of Φ13 was around 50% in the 18 kidneys colonized with a mixed bacterial population (Fig. 1C), a significant increase over the spontaneous excision rate of 0.5% observed during growth *in vitro*. The specific in vivo dynamics of prophage content suggests either an induction of Φ13 excision in response to conditions encountered during infection or a selective advantage for the non-lysogenic strain, potentially due to enhanced fitness during kidney colonization through loss of the prophage.

### Excision of prophage Φ13 is essential for the *S*. *aureus* infectious process

Increased prophage Φ13 excision during infection suggests it may impact virulence. The Φ13 SAOUHSC*-02293* gene, referred to henceforth as *int*, encodes a protein highly similar to tyrosine recombinase integrases such as Int of phage lambda (16, 18). We deleted the entire coding sequence of the Φ13 *int* gene in *S. aureus* strain HG001 in order to prevent excision, locking the prophage within the bacterial chromosome. We also constructed a strain where the entire Φ13 genome is deleted, restoring an intact and functional *hlb* gene (see Materials and Methods). These mutations had no effect on bacterial growth *in vitro* (Supplementary Fig. 2). β-hemolytic activity following Φ13 excision was detected on sheep blood agar plates as a distal cold-inducible halo of partial hemolysis (indicated by the white arrow in Fig. 2A).

**Figure 2:**
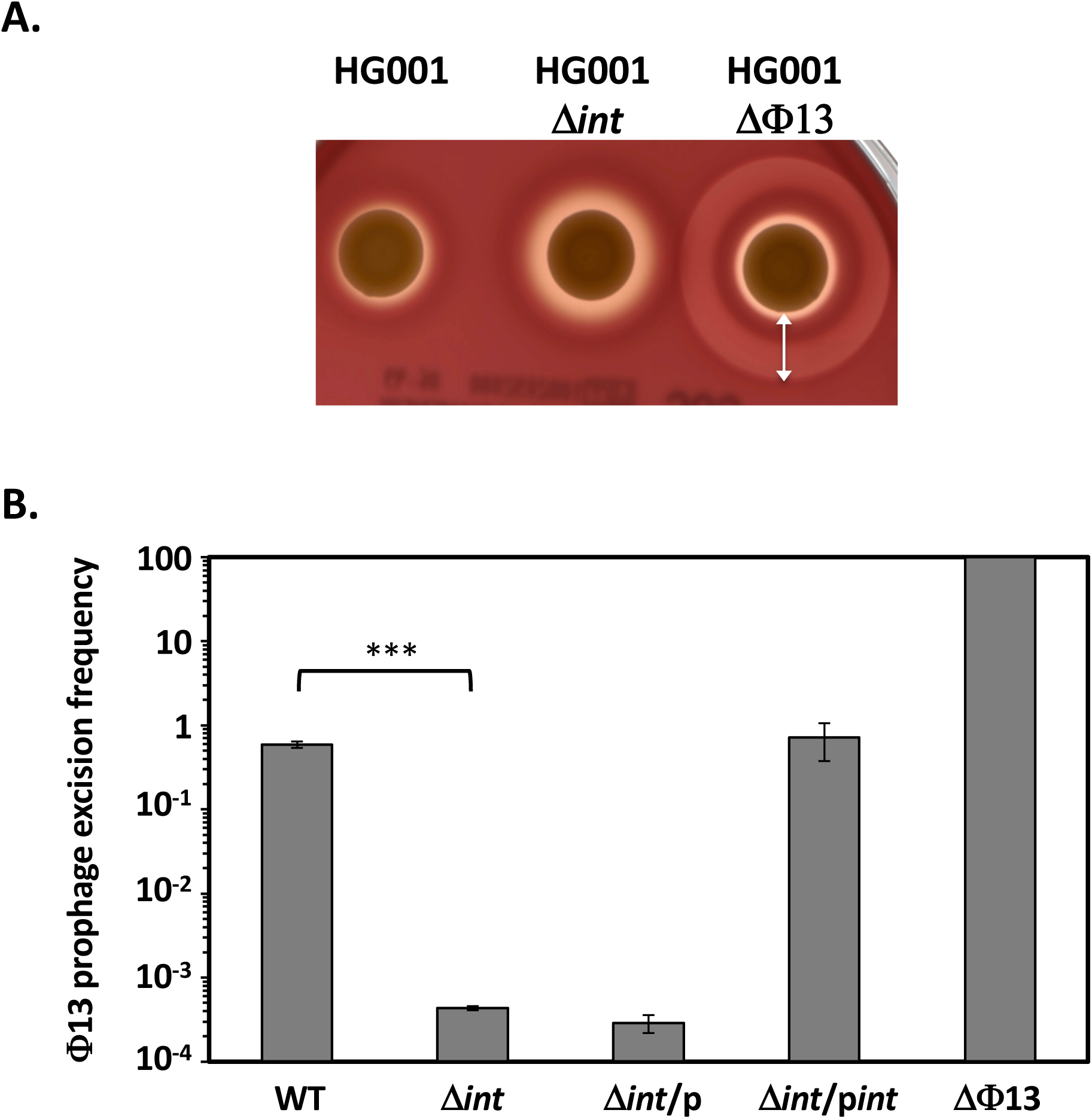
Characterization of the *S. aureus* Δ*int* and ΔΦ**13 mutants.** Strains were grown overnight at 37°C with aeration. A) 3 μl of the bacterial cultures was spotted on sheep blood agar plates and incubated consecutively for 24 h at 37°C and 24 h at 4°C to detect hemolytic activities. The white arrow indicates the β-hemolytic halo characteristic of bacteria lacking prophage Φ13. B) Prophage Φ13 excision frequency detected by qRT-PCR using oligonucleotide pairs OMA7-OMA8 and OMA7-OMA9, specific for the presence or absence of Φ13, respectively. A Φ13 lysogen strain was included as a control to confirm the specificity of the amplicons used to quantify Φ13 lysogeny. Results are presented as means ± SEM of 3 replicates and *t*-test was used to compare WT and Δ*int* results (***, p<0,0001).

As shown in Fig. 2B, the frequency of Φ13 excision in the Δ*int* mutant is lowered more than 1000-fold as compared to that of the wild type strain (0.6%), confirming the essential role of Int in Φ13 excision. Full complementation of the Δ*int* mutant was obtained by expressing the *int* gene from the Pprot constitutive promoter in the pMK4Pprot-*int* plasmid (Fig. 2B).

We next examined the effect on *S. aureus* virulence of either the absence of prophage Φ13 (HG001ΔΦ13 strain, intact *hlb* gene) or the lack of Φ13 excision (HG001Δ*int* strain, *hlb* gene disrupted). As shown in Fig. 3A, using a systemic model of murine infection (5 × 10^7^ c.f.u. intravenous injection), the absence of Φ13 with a resulting intact *hlb* gene led to acute virulence with the rapid death of infected animals (50% mortality 24 h post-infection for the ΔΦ13 strain vs. six days for the wild type strain), and more than 90% mortality at the end of the assay (ten days post-infection).

**Figure 3:**
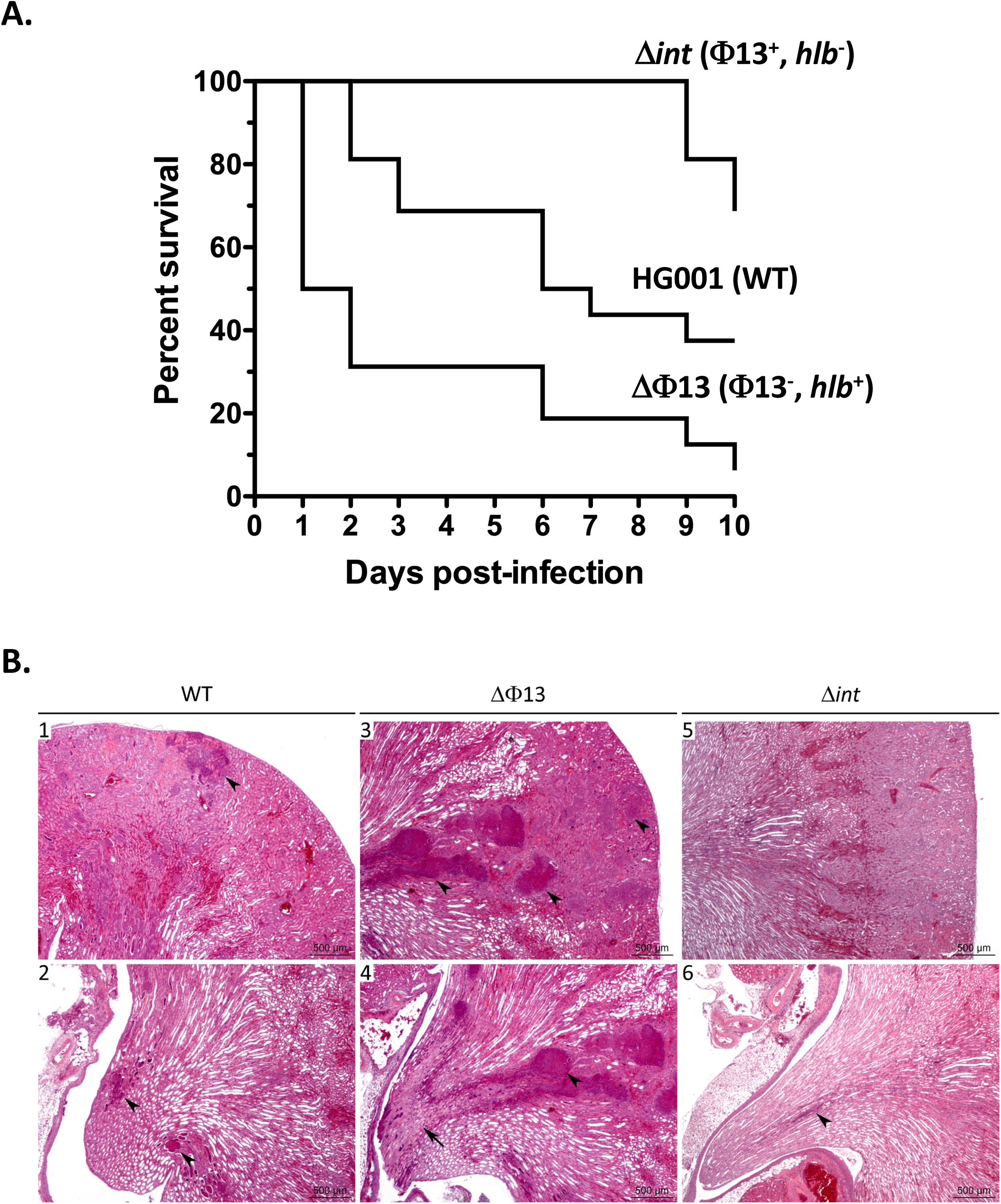
Impact of prophage Φ13 and *hlb* on *S. aureus* virulence. A) Kaplan-Meier survival curves of *RjOrl:SWISS* mice infected intravenously with *S. aureus* HG001 wild type strain and the Δ*int* and ΔΦ13 mutants (5 × 10^7^ c.f.u./injection). A total of 14 mice were used in each group in two independent experiments. For each strain, the genotype is indicated followed by the relevant genetic context (presence or absence of prophage Φ13 and a functional *hlb* gene). Statistical analysis of data using Log-rank (Mantel Cox) test was performed to compare each mutant strain to the HG001 parental strain. WT/Δ*int*: *p*=0.0077; WT/ ΔΦ13 : *p*=0.0021. B) Histopathological kidney analyses 6 days post-infection by intravenous inoculation with sub-lethal bacterial loads (2 × 10^7^ c.f.u./injection) of the *S. aureus* HG001 wild type (1 and 2), Δ*int* (3 and 4) and ΔΦ13 (5 and 6) strains. Most (5/6) mice infected with the wild type strain displayed mild to moderate nephritis, with neutrophilic infiltrates/abscesses (black arrowheads) of varying sizes in the cortex (1), medulla, and papilla (2). Infection with strain ΔΦ13 led to more severe nephritis (5/6 mice), with multifocal abscesses (3 and 4, black arrowheads), extension to tubules and interstitium, and ischemic necrosis of the papilla (4, black arrow). Most mice (4/6) infected with the Δ*int* strain displayed less severe lesions, with mononuclear cell infiltrates most often in the cortex (5, not seen at a low magnification) but also in the papilla (6, arrowhead).

Conversely, when Φ13 excision is abolished (Δ*int* strain), *S. aureus* virulence is strongly reduced compared to the wild type strain (31% mortality at the end of the assay, *vs* 63%). These results indicate that Φ13 excision and its timing are crucial parameters for the *S. aureus* infectious process.

### Φ13 dynamics are involved in organ damage during infection

To understand the effect of Φ13 dynamics on *S. aureus* virulence, we measured bacterial colonization, and evaluated the kidney lesions of infected mice by histopathological analysis. Mice were inoculated intravenously with sub-lethal concentrations of *S. aureus* cells (approximately 2 × 10^7^ c.f.u.). At six days post-infection, mice were sacrificed and one kidney was fixed for histopathological analysis and the other homogenized for c.f.u. counts. Kidney bacterial loads for the parental HG001 strain and the HG001ΔΦ13 and HG001Δ*int* mutants were not significantly different (approximately 5 × 10^7^ c.f.u. per kidney; Supplementary Fig. 3). However, histopathological analysis revealed nephritis (very probably embolic nephritis) with significantly varying degrees of severity between mice infected with the parental HG001 strain or the ΔΦ13 and Δ*int* mutants. At six days post-infection, mice inoculated with the wild type strain generally displayed mild to moderate nephritis (5/6 mice) or multifocal inflammatory infiltrates (macrophages and other mononucleated cells) in the renal cortex (1/6 mouse) (Fig. 3B1 and 3B2). In contrast, the ΔΦ13 strain leads to more severe nephritis (5/6 mice), with neutrophilic inflammation forming multifocal abscesses, extension to tubules and interstitium, and ischemic necrosis of the papillae (Fig. 3B3 and 3B4). Mice infected with the Δ*int* strain displayed less severe lesions, with mononuclear cell infiltrates mostly in the cortex (4/6 mice) or mild to moderate (1/6 mouse) or marked (1/6 mouse) nephritis (Fig. 3B5 and 3B6).

These results show that Φ13 dynamics and correlated β-hemolysin production are not essential for host colonization but instead are key factors in host organ damage during infection.

### Φ13 excision triggers host inflammatory response in infected THP-1 cells

Given the major impact of Φ13 excision on virulence and inflammation, we tested its effect on induction of the cellular inflammatory response using human monocytic THP-1 cells. We compared the bacterial internalization capacity of THP-1 cells, the intracellular viability of bacterial cells and the induction of inflammatory markers following infection with the parental HG001 strain and the ΔΦ13 and Δ*int* mutants.

THP-1 differentiated macrophages were very efficient for *S. aureus* phagocytosis regardless of prophage content, with nearly 100% of bacteria internalized with a m.o.i. (multiplicity of infection) of 10 after 2 hours (Supplementary Fig. 4A). Intracellular survival was not significantly different for the three strains with approximately 10% survival after 24 h post infection and less than 5% after 48 h (Supplementary Fig. 4B). To measure induction of the inflammatory response, duplicate aliquots of infected THP-1 cells were frozen 24 h post-infection, treated to recover cellular RNA and induction of inflammatory cytokine genes measured by qRT-PCR. We measured induction of three pro-inflammatory cytokines previously shown to be induced by *S. aureus* infection, TNFα, IL-1β and IL-6 (19). Although there was no significant difference in induction levels of TNFα and Pro-IL-1β for the three strains, IL-6 gene expression was strongly increased when THP-1 cells were infected with the ΔΦ13 strain as compared to the HG001 strain and the Δ*int* mutant (Fig. 4). Thus, lysogeny by Φ13 and the concomitant loss of β-hemolytic activity appear to down-regulate the inflammatory response through lower induction of the pro-inflammatory cytokine IL-6.

**Figure 4:**
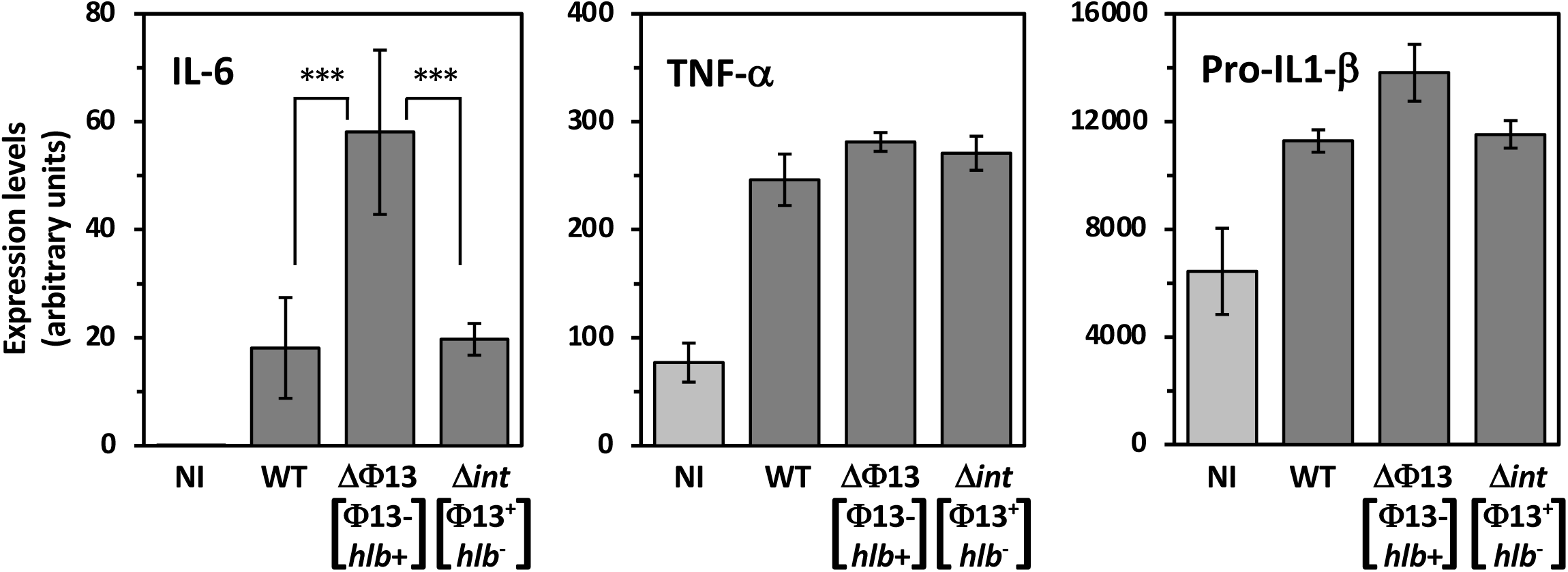
Impact of Φ13 on cytokine induction by THP-1 neutrophils. PMA-activated THP-1 cells were infected with the HG001 wild type strain or the ΔΦ13 and Δ*int* mutants with a m.o.i. = 10. After 24 H of infection, cytokine gene expression was determined by qRT-PCR performed with cDNA synthesized from RNA of THP-1 cells infected by the indicated strains (dark grey) or non-infected control (light grey). Expression levels are given in arbitrary units normalized against that of HPRT. Each experiment was carried out 6 times in two independent series (means ± SD). ***: *P*<0.005 (*t*-test).

### Impact of *hlb* on *S. aureus* virulence

Taken together the results shown above suggest a major role of β-hemolysin production in the increased virulence of the ΔΦ13 strain. To test this hypothesis, we compared the virulence of the ΔΦ13 strain, which produces β-hemolysin, with the virulence of the ΔΦ13Δ*hlb* strain in which the *hlb* gene has also been deleted. As shown in Fig. 5A, 44% of mice infected by the ΔΦ13Δ*hlb* strain survived at nine days post-infection, whereas survival of mice infected by the strain producing β-hemolysin (ΔΦ13) was only 14%, indicating that it is the major factor responsible for the effect of Φ13 excision on *S. aureus* virulence. To test the contribution of Φ13 in virulence, we constructed a strain where both Φ13 and an intact *hlb* gene are present (Δ*int hlb* complemented strain; see Materials and Methods, and Supplementary Table 1). As shown in Fig. 5B, mouse mortality caused by the ΔΦ13 strain producing β-hemolysin but lacking the Φ13 prophage was essentially the same as that following infection by the Δ*int hlb* complemented strain (with both the Φ13 prophage and a functional *hlb* gene). This result confirms the major role of Φ13 dynamics in *S. aureus* virulence by controlling timing of β-hemolysin production during the infectious process.

**Figure 5:**
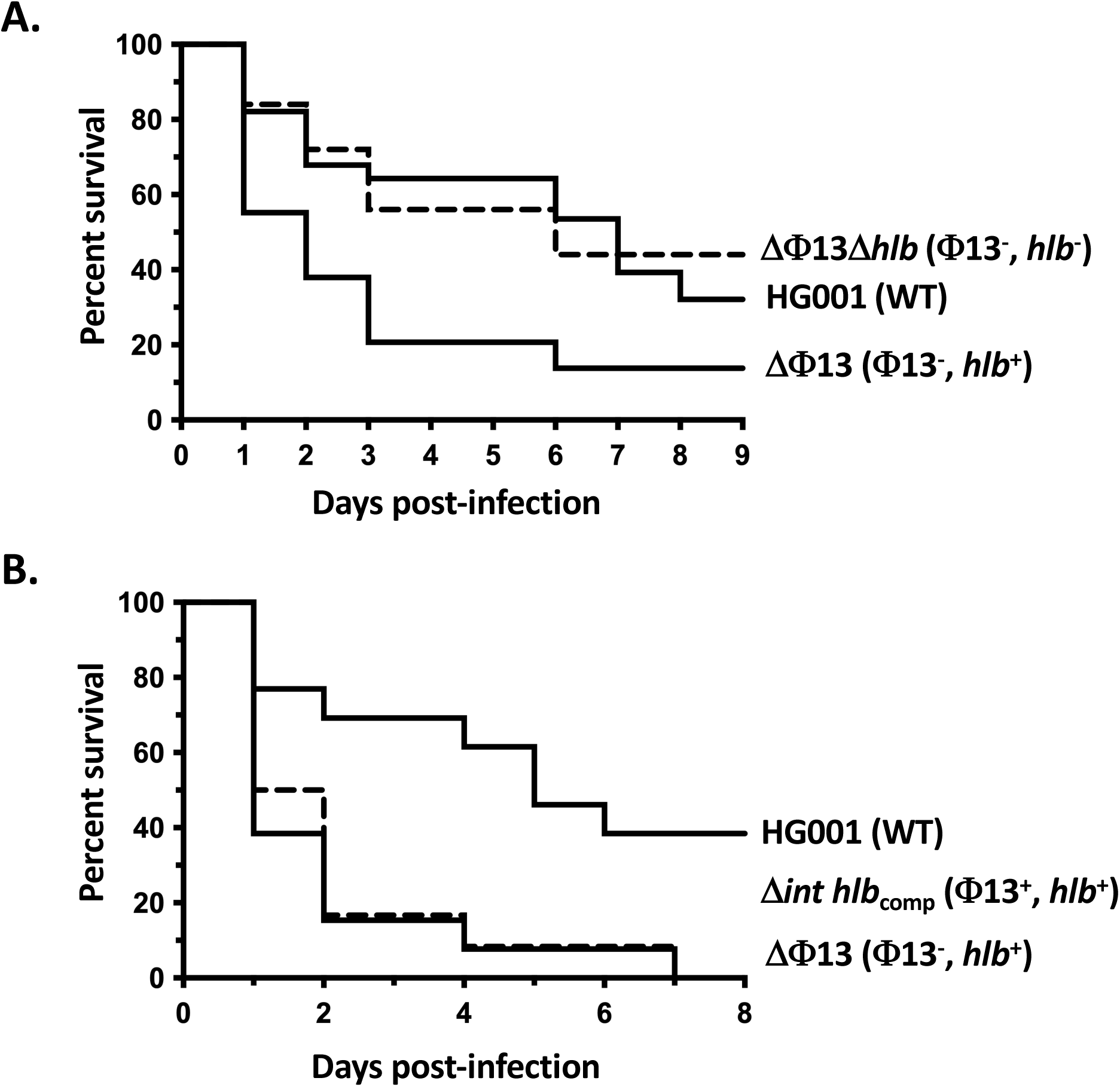
Impact of *hlb* on *S. aureus* virulence. Kaplan-Meier survival curves of *RjOrl:SWISS* mice infected intravenously with *S. aureus* HG001 wild type strain and the Δ*int hlb*_comp_, ΔΦ13 and ΔΦ13Δ*hlb* mutants (5 × 10^7^ c.f.u./injection). A total of 14 mice were used in each group in two independent experiments. *, *P*<0.05 and **, *P*<0.01 (Wilcoxon test). For each strain, the genotype is indicated followed by the relevant genetic context (presence or absence of prophage Φ13 and a functional *hlb* gene). A) Role of *hlb* in virulence by comparing infection with the HG001 strain and the ΔΦ13 and ΔΦ13Δ*hlb* derivatives. Statistical analysis of data using Log-rank (Mantel Cox) test was performed to compare results from each mutant strain to the HG001 parental strain. WT/ΔΦ13Δ*hlb*: *p*=0.6157 (ns); WT/ΔΦ13 : *p*=0.0039. B) Role of Φ13 in virulence by comparing infection with the HG001 strain and the Δ*int hlb*_comp_ (dashed line) and ΔΦ13 derivatives. Statistical analysis of data using Log-rank (Mantel Cox) test was performed to compare data for mutant strains to the HG001 parental strain. WT/Δ*int hlb*_comp_: *p*=0.0147; WT/ΔΦ13 : *p*=0.0066.

## DISCUSSION

*Staphylococcus aureus* is a highly successful human pathogen with a sophisticated arsenal of virulence factors whose production is tightly controlled by multiple regulatory networks, such as two-component systems (20). Further fine-tuning is provided by mobile genetic elements such as pathogenicity islands (SaPI), plasmids, transposons, staphylococcal cassette chromosomes (SCC) and bacteriophages conferring additional virulence factors or antibiotic resistance genes (21). Prophages are widely distributed amongst *S. aureus* strains. Among these, the Φ13 *S. aureus* prophage is inserted within the *hlb* gene encoding β-hemolysin and carries at least three genes involved in host interactions and escape from the immune response, known as the immune evasion cluster (IEC): *sak*, encoding staphylokinase, involved in converting host plasminogen to active plasmin, *scn*, encoding a staphylococcal complement inhibitor, and *chp*, encoding a neutrophil and monocyte chemotaxis inhibitory protein (9). The Φ13 *hlb*-converting prophage family is present in approximately 90% of *S. aureus* clinical isolates (9).

β-hemolysin is a sphingomyelinase that can lyse proliferating human lymphocytes, contributing to the host immune evasion (22). Independently of its pore forming activity, β -hemolysin has a biofilm ligase effect by forming a nucleoprotein matrix both in vitro and during infectious endocarditis (23, 24). It has also been shown that Hlb favors mouse skin colonization (12). We noted that *S. aureus* cells isolated from infected mice frequently displayed β-hemolytic activity due to *in vivo* excision of the Φ13 prophage. This prompted us to examine the effect of phage mobility on host pathogen interactions. Our results indicate that mobility of this prophage is required for full virulence of *S. aureus* as part of the time-course of the infectious process. We identified and inactivated the potential integrase gene of this prophage, effectively locking the prophage within the *S. aureus* genome by preventing its excision, and showed that this leads to strongly decreased virulence, with 70% of the mice still surviving after 10 days. Conversely, a strain lacking the Φ13 prophage exhibited greater virulence than the wild type strain, with 50% of infected mice succumbing within the first day of infection, compared to six days for the wild-type strain. These Φ13-associated virulence traits are not correlated to differences in the strain’s ability to colonize the host, as bacterial loads in the infected kidneys were similar regardless of the strain’s lysogenic status. However, kidney damage following infection was strongly associated with the presence of Φ13, showing a clear gradient where a lower proportion of lysogenic bacteria resulted in more severe tissue lesions. In a human monocyte (THP-1) infection model used to assess inflammatory response, we found that the non-lysogenic strain significantly increased the production of the pro-inflammatory cytokine IL-6. IL-6 triggers the release of acute-phase response proteins into the bloodstream, leading to a systemic inflammatory response (25). It is also considered a key biomarker for predicting poor outcomes in sepsis patients and is a promising target for inhibitors to help mitigate the adverse effects of inflammation (19, 26). The induction of IL-6 by the non-lysogenic strain may be linked to its heightened virulence and the significant neutrophil recruitment observed at the inflammation site, as shown by the histopathological analysis of infected kidneys.

The exact trigger causing excision of Φ13 during infection remains to be established. Interestingly, our observations show that after intravenous infection, kidneys of the same mouse can be colonized differently, one kidney by a lysogenic population, while the other harbors a mixed population. This suggests that Φ13 excision is likely induced in the kidneys following colonization rather than during the initial spread of bacteria through the bloodstream. Interestingly, a previous study using a rabbit infection model found that bacteria producing β-hemolysin emerged during organ colonization, but this was not observed in the bacterial population recovered from the bloodstream (11). Stress conditions activating the SOS response are known to induce prophage excision. However, despite many attempts, we were not able to induce Φ13 excision from the bacterial chromosome through treatment with Mitomycin C as SOS-inducing agent. Similar results were previously described with a Newman strain carrying the closely related *hlb*-converting prophage, ΦNM3 (27). More recently, it has been demonstrated that exposure to oxidative stress (such as hydrogen peroxide) and biofilm growth conditions promote the excision of ΦSA3mw (28). This may correlate with our in vivo observations, i.e., that phage induction occurs after organ colonization, under biofilm development conditions.

Epidemiological studies have shown that *hlb*-converting prophages are more commonly found in strains colonizing humans (from nasal swabs) than in invasive strains (isolated from blood) (3). Additionally, Φ13-derivative carriers are significantly more frequent in human *S. aureus* isolates than in animal strains (29). These *hlb*-converting phages not only inactivate *hlb* but also carry virulence genes, including an immune evasion cluster (IEC) (9). The Sak, SCIN, and CHIPS determinants encoded by this cluster play crucial roles in modulating neutrophil chemotaxis, bacterial phagocytosis, and killing (17). However, their activity is highly specific, primarily targeting the human immune system (30–32). This specificity of the Φ13-derivative IEC in counteracting the human immune response likely contributes to the greater prevalence of *hlb*-converting prophages in human *S. aureus* strains.

Interestingly, we demonstrated using a murine systemic infection model, that the *hlb* gene is the major determinant responsible for the increased virulence observed in the absence of the Φ13 phage. When Hlb is produced, no effect of Φ13 alone was observed (Fig. 5B), likely because the strain is already highly virulent due to constant Hlb production. However, in the absence of *hlb*, the presence of Φ13 led to lowered mouse mortality (Fig. 3A and 5A). While the immune evasion factors encoded by Φ13 have been described as human-specific, it cannot be excluded that they may also exhibit some activity during mouse colonization. Alternatively, Φ13 may encode uncharacterized proteins that could modulate *S. aureus* virulence.

Excision of Φ13 and the chromosomal rearrangement reconstituting an intact *hlb* gene can be viewed as a timing device, ensuring that the bacteria first colonize the host, benefiting from the immune evasion cluster proteins encoded by the prophage, with production of β-hemolysin and host tissue damage occurring once the bacteria have reached a threshold level, in agreement with the temporal control of *Staphylococcus aureus* virulence factors (33). One of the first examples of prophage excision as a timing device for a developmental process is the *skin* element whose excision leads to a chromosomal rearrangement reconstituting the gene encoding the Sigma K sigma factor, essential for *Bacillus subtilis* sporulation (34, 35).

As demonstrated in this study and supported by the literature, the heterogeneity within *S. aureus* populations with respect to *hlb*-converting prophages appears to be a common trait. This could be seen as a bet-hedging strategy, allowing the bacteria to retain the ability to produce both immune evasion proteins (in the lysogenic population) and the potent Hlb cytotoxin (in the non-lysogenic cells).

Additionally, since both β-hemolysin and the immune evasion cluster proteins encoded by the prophage are secreted, it is plausible that functional complementation may occur within a mixed population of lysogenic and non-lysogenic bacteria. Similar dual strategies have been observed before, where bistability leads to cell-to-cell variation in gene expression (36, 37). This mixed population with distinct genetic content would be well-equipped to resist host defenses during colonization as well as causing cellular damage.

## MATERIALS AND METHODS

### Bacterial strains and growth media

*Escherichia coli* K12 strain DH5α™ (Invitrogen) was used for cloning experiments. *Staphylococcus aureus* strain HG001, derived from strain NCTC 8325 by repairing the mutated *rsbU* gene (15), was used for genetic and functional studies. Plasmids were first transformed into the restriction deficient *S. aureus* strain RN4220 before introduction into the HG001 or derivative strains. *E. coli* strains were grown in LB medium with ampicillin (100 μg/mL) added when required. *S. aureus* strains and plasmids used in this study are listed in Supplementary Table 1. *S*. *aureus* strains were grown in Trypticase Soy Broth (TSB; Difco) supplemented with chloramphenicol (10 μg/mL) or erythromycin (1 μg/mL) when required. *E*. *coli* and *S*. *aureus* strains were transformed by electroporation using standard protocols (38) and transformants were selected on LB or Trypticase Soy Agar (TSA; Difco) plates, respectively, with the appropriate antibiotics.

### DNA manipulations

Oligonucleotides used in this study were synthesized by Eurofins Genomics and are listed in Supplementary Table 2. *S*. *aureus* chromosomal DNA was isolated using the MasterPure^TM^ Gram-positive DNA purification Kit (Epicentre Biotechnologies). Plasmid DNA was isolated using a QIAprep Spin Miniprep kit (Qiagen) and PCR fragments were purified using the Qiaquick PCR purification kit (Qiagen). T4 DNA ligase and restriction enzymes, PCR reagents and Q5 high-fidelity DNA polymerase (New England Biolabs) were used according to the manufacturer’s recommendations. Nucleotide sequencing of plasmid constructs was carried out by Eurofins Genomics .

### Plasmids and mutant strain construction

Construction of the ΔΦ13 mutant strain was carried out by cloning an 1800 bp DNA fragment centered around the *attB* site, generated by PCR using oligonucleotides OSA418 and OSA419 and chromosomal DNA from the SH1000 strain (a NCTC8325-4 derivative cured for prophages), between the *Bam*HI and *Eco*RI sites of the pMAD vector (39), to give plasmid pMAD-Φ13. For deletion of the Φ13 integrase (*int*; *saouhsc-02239*), DNA fragments corresponding to the *int* upstream and downstream regions were generated using oligonucleotides OSA411/OSA412 (900 bp) and OSA413/OSA414 (820 bp) cloned between the pMAD *Bam*HI and *Nco*I sites. The *aad*(9) spectinomycin resistance gene (40) was inserted in the center of the 23 nt intergenic region between the convergent Φ13 *saouhsc-02232* and *saouhsc-02233* genes, to avoid any polar effects on transcription. DNA fragments corresponding to the regions upstream and downstream from the insertion site were generated by PCR using primers OSA508/OSA509 and OSA510/OSA511, respectively, and primers OSA299/OSA470 for the *aad*(9) cassette with plasmid pIC333 (41) as the DNA matrix.

The DNA fragments were cloned into pMAD using the *Bam*HI/*Sal*I sites for the upstream fragment, *Sal*I/*Nco*I for the *aad*(9) cassette, and the *Nco*I/*Bgl*II sites for the downstream fragment, generating plasmid pMAD-Φ13*aad*(9). For deletion of *hlb*, we generated upstream and downstream fragments by PCR using oligonucleotides OSA442/OSA443 (884 bp) and OSA444/OSA445 (889 bp) respectively. These fragments were introduced into pMAD between the *Bam*HI and *Eco*RI sites, thus generating the pMAD-*hlb* plasmid. To introduce a functional *hlb* allele into the Φ13 lysogenic strain, a pMAD-*hlb*comp plasmid was constructed to place the intact *hlb* gene upstream from the Φ13 genome. A PCR fragment including the *hlb* upstream region and the *hlb* intact gene (1760 bp) was generated using the OSA418 and OSA573 primers and the SH1000 genomic DNA as matrix and cloned into pMAD using the *BamH*I and *Sal*I restriction sites. A second PCR fragment corresponding to the part of Φ13 adjacent to *att*L was generated with the OSA574-575 primers and cloned between the *Sal*I and *Bgl*II restriction sites of pMAD.

Plasmids were introduced by electroporation into *S*. *aureus* strain RN4220, and transformants selected at 30°C on TSA plates containing erythromycin and 5-bromo-4-chloro-3-indolyl-β-D-galactopyranoside (X-Gal) (100 µg/mL). Plasmids were then introduced into strain HG001 and integration and excision performed as previously described (39), yielding strains HG001ΔΦ13, HG001Δ*int*, and HG001Φ13-aad(9) and into strain HG001ΔΦ13 for pMAD-*hlb* and HG001Δ*int* for pMAD-*hlb*comp to give strains HG001Δ*int hlb*comp and HG001ΔΦ13Δ*hlb*.

Plasmid pMK4Pprot-*int* was used for complementation of the Δ*int* mutant and constructed by cloning the entire *int* coding sequence (amplicon OSA446/OSA447) under the control of the constitutive P*prot* promoter (42).

### Prophage content measurement by quantitative real time PCRs (qRT-PCRs)

Strains were grown overnight in TSB at 37°C, with aeration. Cells were pelleted by centrifugation (2 min, 20,800 *g*) from 2 mL culture samples and genomic DNA extracted using the MasterPure^TM^ Gram Positive DNA Purification Kit (Tebu-Bio).

Oligonucleotides were designed with the BEACON Designer 4.02 software (Premier Biosoft International, Palo Alto, CA) to synthesize 100-200 bp amplicons (see Supplementary Table 2). Quantitative real-time PCRs (qRT-PCRs) and critical threshold cycles (CT) were performed and determined using the SsoFast^TM^ EvaGreen Supermix (Bio-Rad, Hercules, CA). All assays were performed using quadruplicate technical replicates, and repeated with three independent biological samples, with one representative experiment shown. Prophage loss frequency was calculated using the formula: 1/(1+2^ΔCt^), ΔCt= Ct_phage negative_-Ct_phage positive_ derived from the ΔCt method (43).

### Hemolysis detection

Bacteria were grown overnight in TSB at 37°C with aeration. Culture aliquots (3 μl) were spotted on Columbia plates containing 5% sheep blood (COS, Biomerieux Craponne). Plates were incubated for 24h at 37°C and transferred to 4°C for an additional 24h to for detection of β-hemolysis (44).

### THP-1 infection experiments

THP-1 monocytes were maintained as previously described (45). Differentiation and adherence were obtained by incubation with Phorbol 12-Myristate 13-Acetate (PMA, 50 ng/mL) during 4 days at 37°C in a 5%CO_2_ atmosphere. Bacterial uptake experiments and survival assays were carried out as previously described (46). Briefly, bacteria were grown in TSB until OD_600nm_ = 1. Cultures were washed in PBS and adjusted to the desired inoculum in RPMI 1640 medium (Gibco), and c.f.u. counts determined by plating serial dilutions on TSA plates. Differentiated THP-1 macrophages were incubated with bacteria (m.o.i. **=** 10) in RPMI 1640 at 37**°**C with 5% CO2 for 1 h to allow bacterial phagocytosis. They were then washed with RPMI and incubated in RPMI-10% Fetal Calf Serum with streptomycin (100 μg/mL), penicillin (100 U/mL), and gentamycin 40 μg/mL to eliminate extracellular bacteria. At the

indicated times, infected macrophages were washed once with RPMI and lysed by incubation in ice-cold water for 15 min and c.f.u. counts were determined by plating serial dilutions on TSA plates. For cytokine gene expression studies by qRT-PCR, THP-1 cells were frozen at -20°C 24h post-infection. Total RNA was extracted using a RNeasy mini kit (Qiagen, CA, USA). Cytokine gene expression was analysed by qRT-PCR using the StepOnePlus real-time PCR system (Applied Biosystems). Primers used for qRT-PCR were TNFα, IL6, pro-IL1beta and HPRT as the reference gene. The qRT-PCRs were performed by Taqman and results presented as relative quantification of gene expression using the 2^−ΔΔCT^ method.

### Animal experiments

Seven-week-old female *RjOrl:SWISS* mice (Janvier Labs) were inoculated intravenously with the specified *S. aureus* strains. To analyze Φ13 excision during the infectious process, mice were infected with a sublethal dose (2 × 10^7^ c.f.u.) of the Φ13-*aad*(9) strain. Mice were sacrificed four days post-infection and kidneys harvested and homogenized for c.f.u. determination following dilution and plating on TSA and TSA with spectinomycin (100 μg/mL). The rate of prophage excision was calculated as [(c.f.u. on TSA) - (c.f.u. on TSA+spectinomycin)]/(c.f.u. on TSA).

Strain HG001 and the Δ*int*, ΔΦ*13*, ΔΦ*13*Δ*hlb* and Δ*int hlb*comp mutants were used for survival studies. Groups of seven mice were injected in the caudal lateral vein with 5 × 10^7^ c.f.u. per mouse in 0.2 mL. Survival was monitored daily for 9 days following infection and three independent experiments were carried out. To measure kidney colonization and histopathological analysis, mice were infected intravenously with a sublethal dose (1 × 10^7^ c.f.u.) of the HG001 parental and Δ*int* and ΔΦ*13* mutant strains (six mice for each strain). At six days post-infection, kidneys were collected and for each mouse, bacterial c.f.u. counts were carried out using one kidney and histopathological observations were done on the other kidney. For histopathology, kidneys were removed and immediately fixed in 10% neutral buffered formalin. After 48 h of fixation, the kidneys were longitudinally sectioned through the hilus in order to obtain the renal cortex, medulla, crest, and pelvis on the same section. These samples were embedded in paraffin, and 4-μm sections were cut and stained with hematoxylin and eosin (lesion description) and Gram (bacterial detection).

### Ethics statement

Animal experiments were carried out at the Institut Pasteur according to European Union guidelines for handling of laboratory animals (http://ec.europa.eu/environment/chemicals/lab_animals/index_en.htm). Animals were monitored daily with a particular attention for any signs of distress. Protocols used were approved by the Institut Pasteur ethics committee (dap160057).

### Statistical analysis

Statistical significance in survival experiments was determined using the log-rank, Mantel–Cox test, and significance across means was carried out using *t*-test (Graph-Pad Prism Software).

## ACKNOWLEDGMENTS

We are grateful to Grégory Jouvion for the histopathological analysis, Patrick Trieu-Cuot, in whose lab this work was carried out, and Shaynoor Dramsi for helpful discussion. All authors have read and agreed to the submitted version of the manuscript.

## FUNDING

This work was supported by funds from Institut Pasteur, CNRS, and Vaincre La Mucoviscidose Association (Grant «Phage induction, a novel essential factor for *Staphylococcus aureus* infections?»).

## AUTHOR CONTRIBUTIONS

Conceptualization: Sarah Dubrac, Lhousseine Touqui, Tarek Msadek.

Data curation: Olivier Poupel, Sarah Dubrac, Lhousseine Touqui, Tarek Msadek.

Formal analysis: Sarah Dubrac, Lhousseine Touqui, Tarek Msadek.

Funding acquisition: Sarah Dubrac, Lhousseine Touqui, Tarek Msadek.

Investigation: Olivier Poupel, Gérard Kénanian, Charlotte Abrial, Sarah Dubrac.

Methodology: Olivier Poupel, Sarah Dubrac.

Project administration: Tarek Msadek, Sarah Dubrac.

Resources: Tarek Msadek, Sarah Dubrac.

Supervision: Tarek Msadek, Sarah Dubrac.

Validation: Tarek Msadek, Sarah Dubrac.

Writing – original draft: Tarek Msadek, Sarah Dubrac.

Writing – review & editing: Tarek Msadek, Sarah Dubrac.

## ADDITIONAL FILES

The following material is available online.

Supplemental Material

**Supplemental Figures and Tables.** Fig. S1–S4 and Tables S1–S2.

## Supplementary Information

**Supplementary Data Fig. 1.**
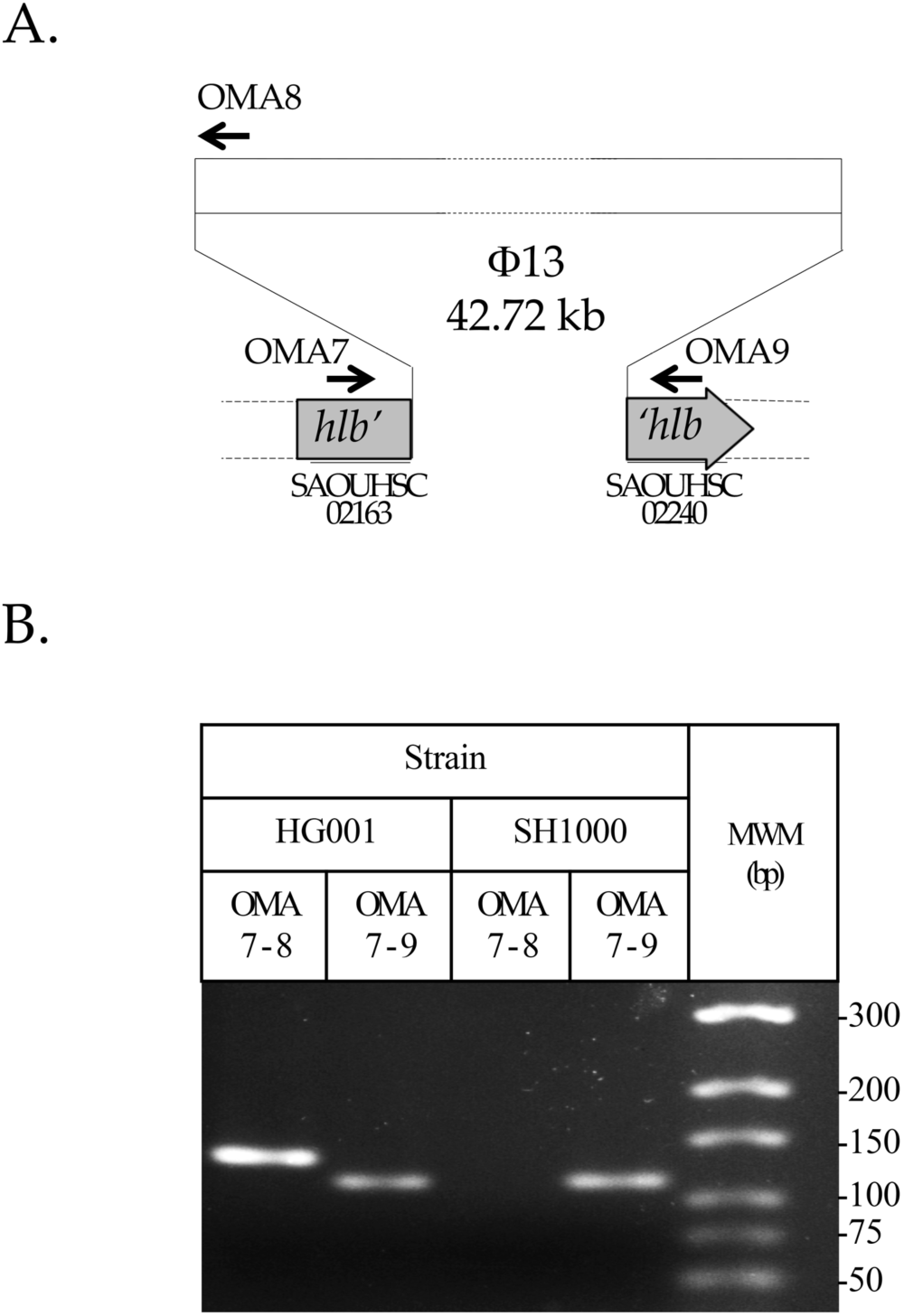
PCR-based detection of phage Φ13 insertion and excision. A: The Φ13 chromosomal insertion locus in strain HG001 (not to scale). The Φ13 *att*P sequence is homologous to the *att*B site located within the *hlb* gene encoding β-hemolysin. Φ13 insertion thus results in disruption and inactivation of the *hlb* gene. Specific primers used to quantify the proportion of lysogenic bacteria are indicated, pair OMA7-OMA8 is specific for Φ13 insertion and pair OMA7-OMA9 for the absence of Φ13. B: Specific amplicons obtained by PCR with the OMA7-8 and OMA7-9 primer pairs using genomic DNA from either the SH1000 strain, cured for Φ13 (114 bp), or from the Φ13 lysogen strain HG001 (132 and 114 bp).

**Supplementary Data Fig. 2.**
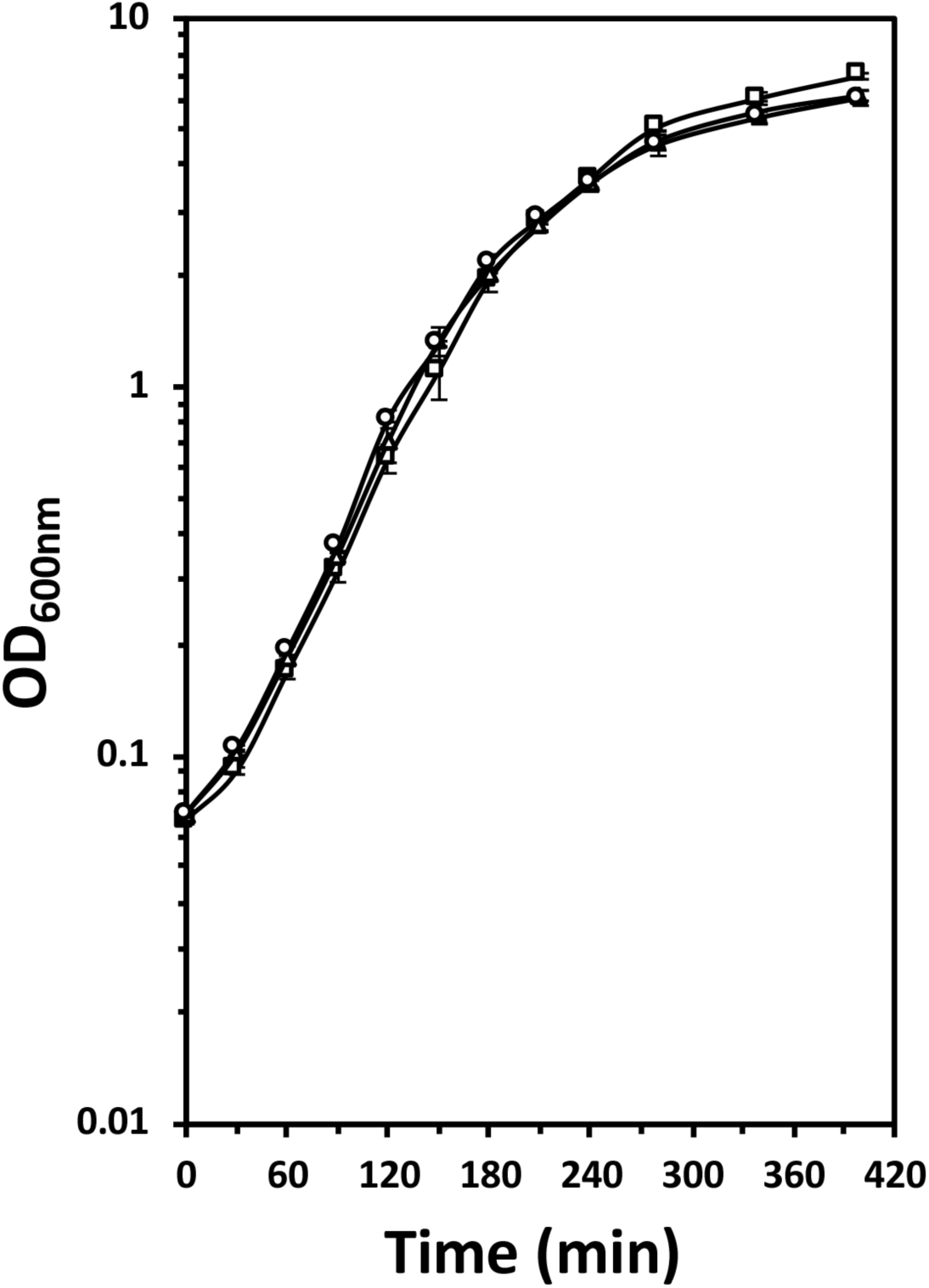
Growth curves of *S*. *aureus* strains HG001 and the ΔΦ13 and Δ*int* mutants. Bacterial cultures were grown overnight, inoculated in TSB at a calculated OD_600nm_ of 0.05 and incubated at 37°C with aeration. Optical densities were followed over a 6.5 hour period. Results are shown as the mean and standard deviation of three independent growth curves. Doubling times (http://www.doubling-time.com/compute.php) were calculated during the exponential growth phase (between 60 min and 150 min) and gave values of approximately 32 min for each strain. Strains: HG001 (□); ΔΦ13 (△); Δ*int* (O).

**Supplementary Data Fig. 3.**
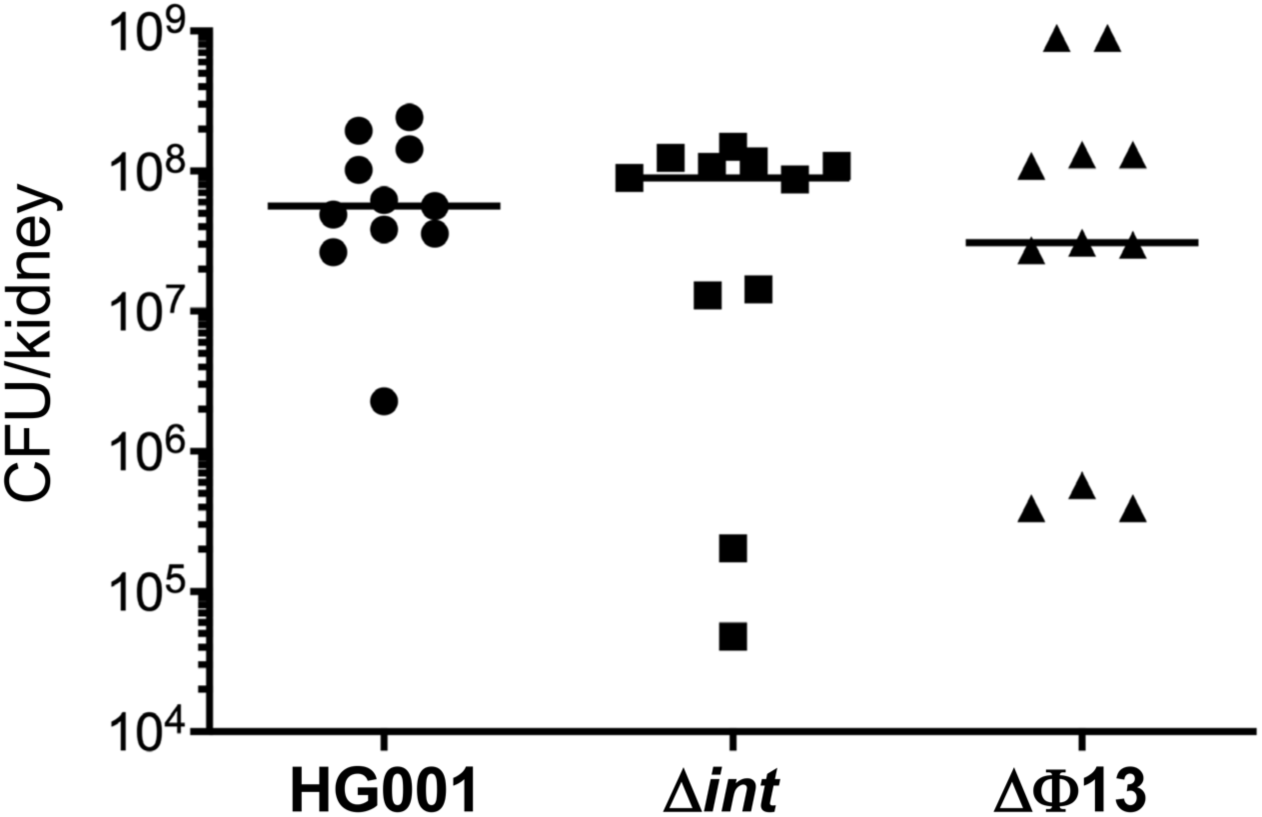
**Kidney colonization following infection by *S. aureus*** *RjOrl:SWISS* mice were infected with the *S. aureus* HG001 wild type strain or the Δ*int* and ΔΦ13 derivatives by intravenous injections with a sub-lethal bacterial load (2.10^7^cfu/injection). Six days post-infection, mice were sacrificed, kidneys were harvested and CFU counts carried out. A total of 11 mice were studied in two independent experiments and the data are compilation of the bacterial counts.

**Supplementary Data Fig. 4.**
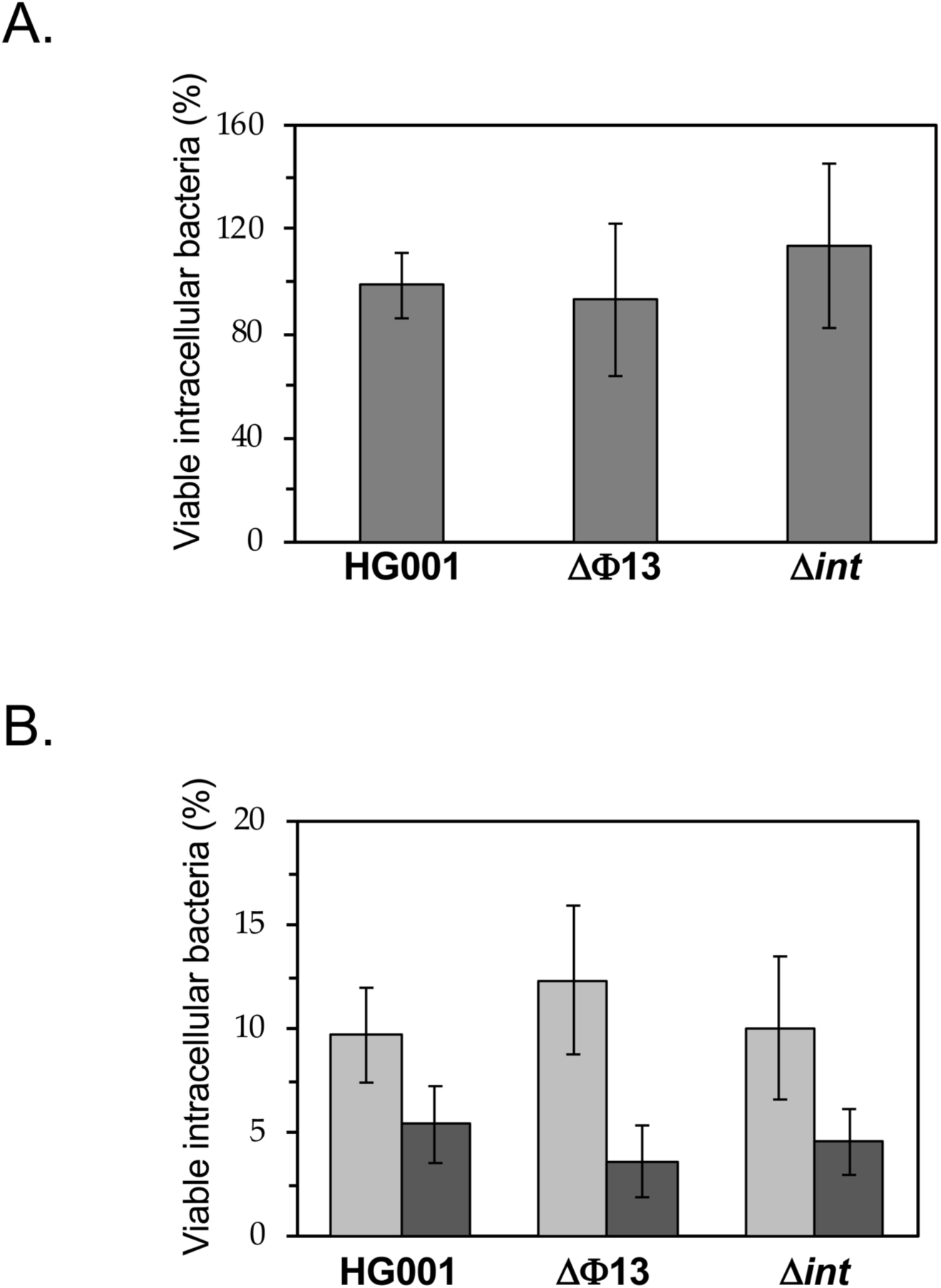
Impact of Φ13 on internalization and survival within THP-1 neutrophils. PMA-activated THP-1 cells were infected with either the HG001 wild type strain or the ΔΦ13 and Δ*int* mutants with a moi=10. A: Viable intracellular bacteria were quantified two hours post-infection to determine the efficiency of internalization. B: Viable bacteria were quantified 24 H (light grey) and 48 H (dark grey) post-infection to determine the capacity of each strain to survive within neutrophils. Results are presented as means and SEM of 3 biological replicates.

**Supplementary Data Table 1:**
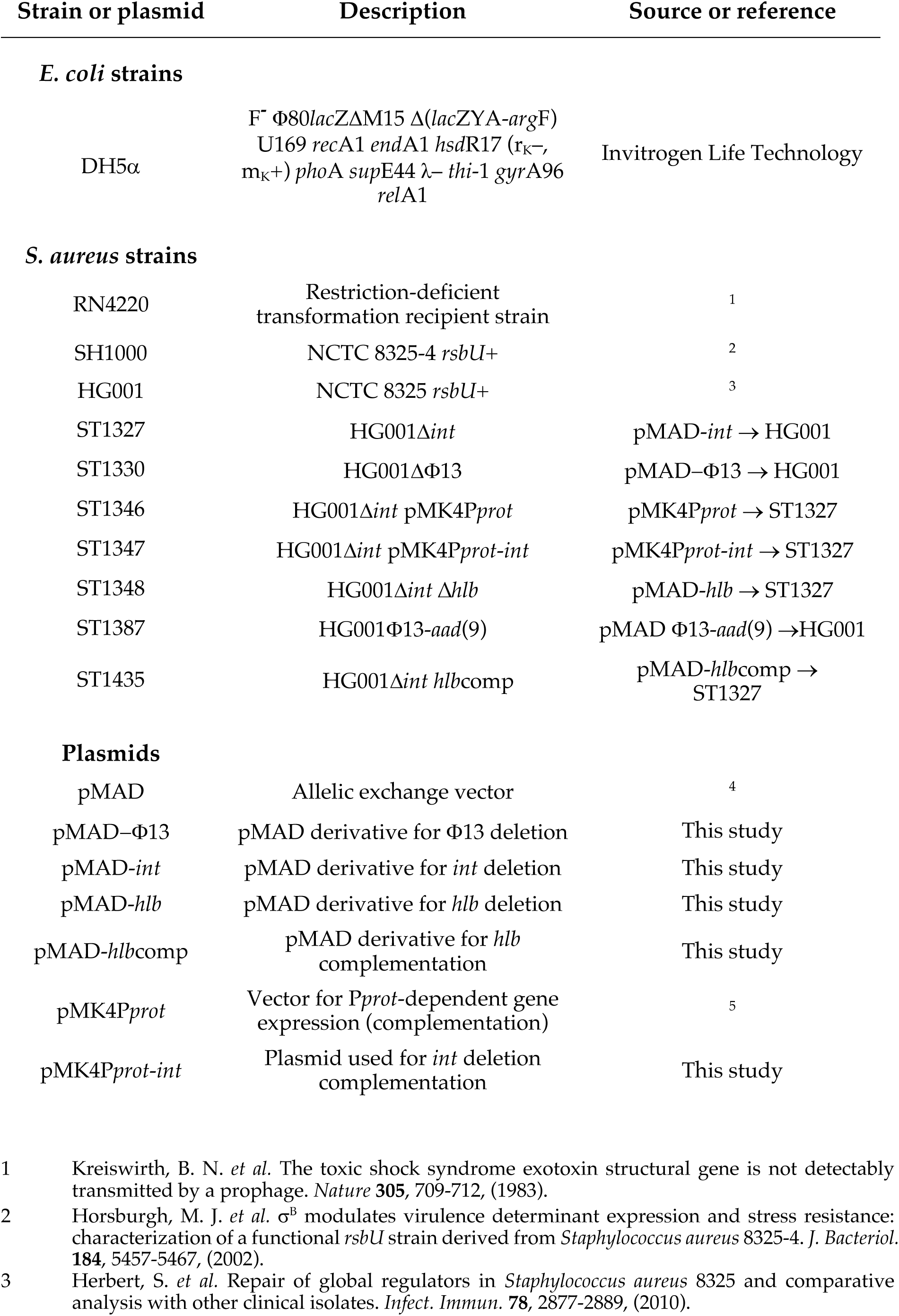
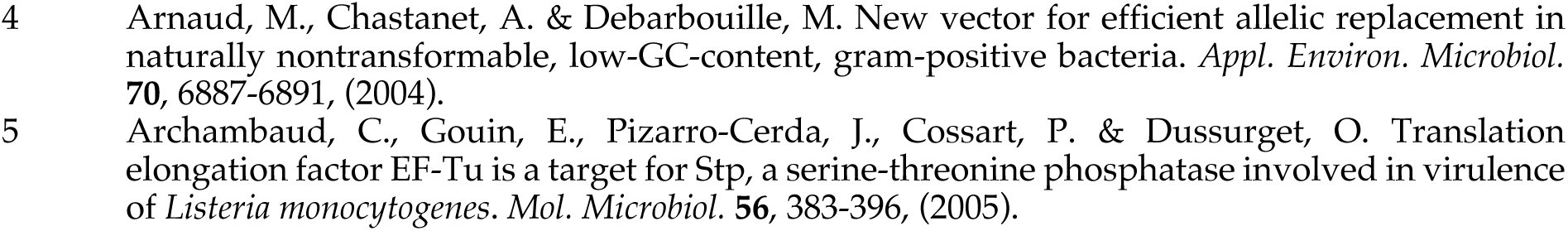
Strains and plasmids used in this study.

**Supplementary Data Table 2:**
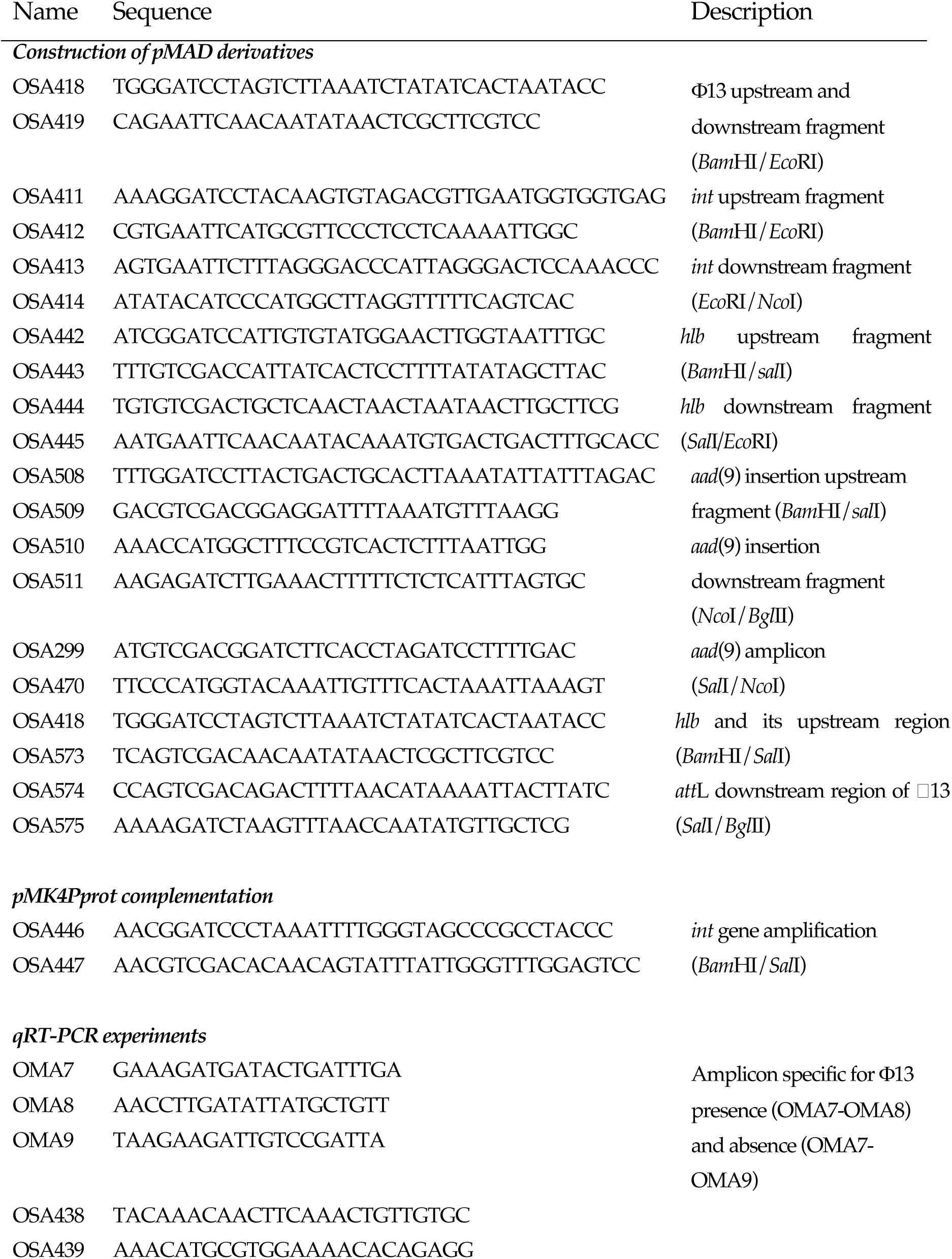
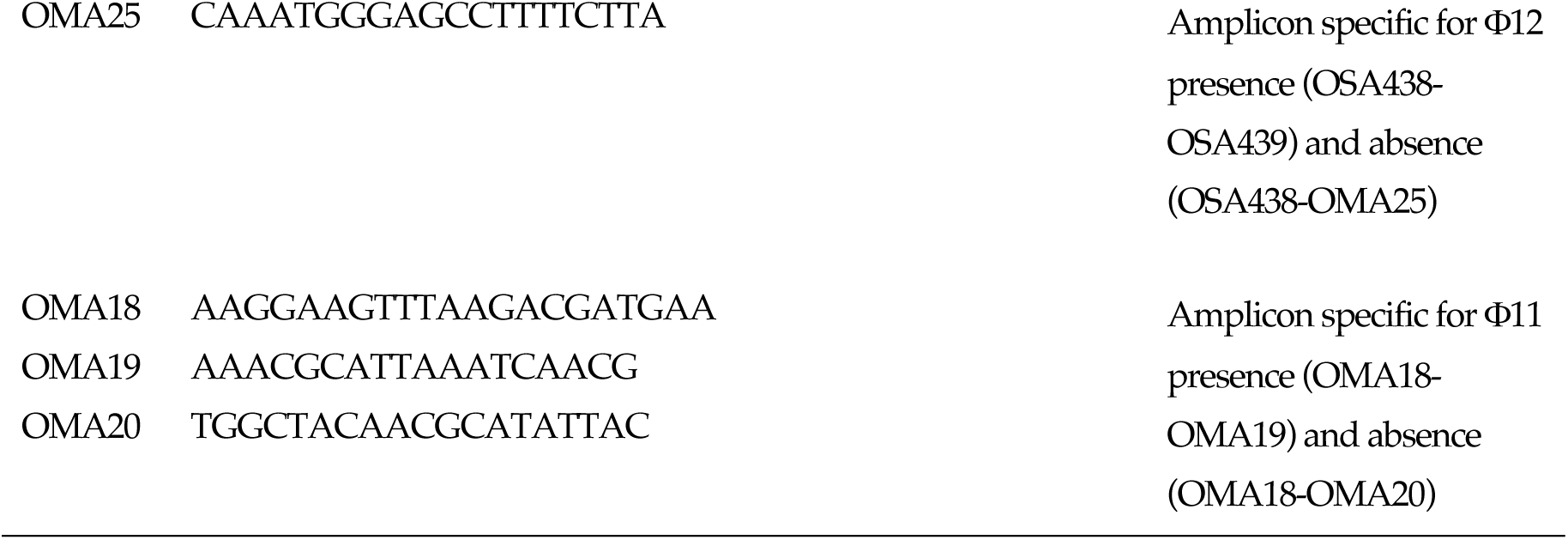
Oligonucleotides used in this study.

## Notes

### Competing Interest Statement

The authors have declared no competing interest.

### Summary of Updates

Title change and multiple minor text modifications, New Supplementary Fig S1B

## REFERENCES

1. Piewngam P, Otto M. 2024. *Staphylococcus aureus* colonisation and strategies for decolonisation. Lancet Microbe 5:e606–e618.

2. Fitzgerald JR, Sturdevant DE, Mackie SM, Gill SR, Musser JM. 2001. Evolutionary genomics of *Staphylococcus aureus*: insights into the origin of methicillin-resistant strains and the toxic shock syndrome epidemic. Proc Natl Acad Sci USA 98:8821–6.

3. Goerke C, Pantucek R, Holtfreter S, Schulte B, Zink M, Grumann D, Broker BM, Doskar J, Wolz C. 2009. Diversity of prophages in dominant *Staphylococcus aureus* clonal lineages. J Bacteriol 191:3462–8.

4. Xia G, Wolz C. 2014. Phages of *Staphylococcus aureus* and their impact on host evolution. Infect Genet Evol 21:593–601.

5. Goerke C, Wirtz C, Fluckiger U, Wolz C. 2006. Extensive phage dynamics in *Staphylococcus aureus* contributes to adaptation to the human host during infection. Mol Microbiol 61:1673–85.

6. Feiner R, Argov T, Rabinovich L, Sigal N, Borovok I, Herskovits AA. 2015. A new perspective on lysogeny: prophages as active regulatory switches of bacteria. Nat Rev Microbiol 13:641–50.

7. Iandolo JJ, Worrell V, Groicher KH, Qian Y, Tian R, Kenton S, Dorman A, Ji H, Lin S, Loh P, Qi S, Zhu H, Roe BA. 2002. Comparative analysis of the genomes of the temperate bacteriophages phi 11, phi 12 and phi 13 of *Staphylococcus aureus* 8325. Gene 289:109–18.

8. Coleman DC, Sullivan DJ, Russell RJ, Arbuthnott JP, Carey BF, Pomeroy HM. 1989. *Staphylococcus aureus* bacteriophages mediating the simultaneous lysogenic conversion of beta-lysin, staphylokinase and enterotoxin A: molecular mechanism of triple conversion. J Gen Microbiol 135:1679–97.

9. van Wamel WJ, Rooijakkers SH, Ruyken M, van Kessel KP, van Strijp JA. 2006. The innate immune modulators staphylococcal complement inhibitor and chemotaxis inhibitory protein of *Staphylococcus aureus* are located on beta-hemolysin-converting bacteriophages. J Bacteriol 188:1310–5.

10. Rohmer C, Wolz C. 2021. The role of hlb-converting bacteriophages in *Staphylococcus aureus* host adaption. Microb Physiol 31:109–122.

11. Salgado-Pabon W, Herrera A, Vu BG, Stach CS, Merriman JA, Spaulding AR, Schlievert PM. 2014. *Staphylococcus aureus* beta-toxin production is common in strains with the beta-toxin gene inactivated by bacteriophage. J Infect Dis 146:784–92.

12. Katayama Y, Baba T, Sekine M, Fukuda M, Hiramatsu K. 2013. Beta-hemolysin promotes skin colonization by *Staphylococcus aureus*. J Bacteriol 195:1194–203.

13. Goerke C, Matias y Papenberg S, Dasbach S, Dietz K, Ziebach R, Kahl BC, Wolz C. 2004. Increased frequency of genomic alterations in *Staphylococcus aureus* during chronic infection is in part due to phage mobilization. J Infect Dis 189:724–34.

14. Hedstrom SA, Malmqvist T. 1982. Sphingomyelinase activity of *Staphylococcus aureus* strains from recurrent furunculosis and other infections. Acta Pathol Microbiol Immunol Scand B 90:217–20.

15. Herbert S, Ziebandt AK, Ohlsen K, Schafer T, Hecker M, Albrecht D, Novick R, Gotz F. 2010. Repair of global regulators in *Staphylococcus aureus* 8325 and comparative analysis with other clinical isolates. Infect Immun 78:2877–89.

16. Gillaspy AF, Worrell V, Orvis J, Roe BA, Dyer DW, Iandolo JJ. 2006. The *Staphylococcus aureus* NCTC 8325 Genome, p 381-412. *In* Fischetti VA, Novick RP, Ferretti JJ, Portnoy DA, Rood JI (ed), Gram-Positive Pathogens, 2nd ed. ASM Press, Washington, DC.

17. Rooijakkers SH, van Kessel KP, van Strijp JA. 2005. Staphylococcal innate immune evasion. Trends Microbiol 13:596–601.

18. Nunes-Duby SE, Kwon HJ, Tirumalai RS, Ellenberger T, Landy A. 1998. Similarities and differences among 105 members of the Int family of site-specific recombinases. Nucleic Acids Res 26:391–406.

19. van den Berg S, Laman JD, Boon L, ten Kate MT, de Knegt GJ, Verdijk RM, Verbrugh HA, Nouwen JL, Bakker-Woudenberg IA. 2013. Distinctive cytokines as biomarkers predicting fatal outcome of severe *Staphylococcus aureus* bacteremia in mice. PLoS One 8:e59107.

20. Haag AF, Bagnoli F. 2017. The role of two-component signal transduction systems in *Staphylococcus aureus* virulence regulation. Curr Top Microbiol Immunol 409:145–198.

21. Lindsay JA. 2010. Genomic variation and evolution of *Staphylococcus aureus*. Int J Med Microbiol 300:98–103.

22. Huseby M, Shi K, Brown CK, Digre J, Mengistu F, Seo KS, Bohach GA, Schlievert PM, Ohlendorf DH, Earhart CA. 2007. Structure and biological activities of beta toxin from *Staphylococcus aureus*. J Bacteriol 189:8719–26.

23. Herrera A, Vu BG, Stach CS, Merriman JA, Horswill AR, Salgado-Pabon W, Schlievert PM. 2016. *Staphylococcus aureus* beta-toxin mutants are defective in biofilm ligase and sphingomyelinase activity, and causation of infective endocarditis and sepsis. Biochemistry 55:2510–7.

24. Huseby MJ, Kruse AC, Digre J, Kohler PL, Vocke JA, Mann EE, Bayles KW, Bohach GA, Schlievert PM, Ohlendorf DH, Earhart CA. 2010. Beta toxin catalyzes formation of nucleoprotein matrix in staphylococcal biofilms. Proc Natl Acad Sci USA 107:14407–12.

25. Tanaka T, Narazaki M, Kishimoto T. 2014. IL-6 in inflammation, immunity, and disease. Cold Spring Harb Perspect Biol 6:a016295.

26. Cao YZ, Tu YY, Chen X, Wang BL, Zhong YX, Liu MH. 2012. Protective effect of Ulinastatin against murine models of sepsis: inhibition of TNF-alpha and IL-6 and augmentation of IL-10 and IL-13. Exp Toxicol Pathol 64:543–7.

27. Bae T, Baba T, Hiramatsu K, Schneewind O. 2006. Prophages of *Staphylococcus aureus* Newman and their contribution to virulence. Mol Microbiol 62:1035–47.

28. Tran PM, Feiss M, Kinney KJ, Salgado-Pabon W. 2019. ∅Sa3mw prophage as a molecular regulatory switch of *Staphylococcus aureus* β-toxin production. J Bacteriol 201:e00766–18.

29. Verkaik NJ, Benard M, Boelens HA, de Vogel CP, Nouwen JL, Verbrugh HA, Melles DC, van Belkum A, van Wamel WJ. 2011. Immune evasion cluster-positive bacteriophages are highly prevalent among human *Staphylococcus aureus* strains, but they are not essential in the first stages of nasal colonization. Clin Microbiol Infect 17:343–8.

30. de Haas CJ, Veldkamp KE, Peschel A, Weerkamp F, Van Wamel WJ, Heezius EC, Poppelier MJ, Van Kessel KP, van Strijp JA. 2004. Chemotaxis inhibitory protein of *Staphylococcus aureus*, a bacterial antiinflammatory agent. J Exp Med 199:687–95.

31. Kwiecinski J, Josefsson E, Mitchell J, Higgins J, Magnusson M, Foster T, Jin T, Bokarewa M. 2010. Activation of plasminogen by staphylokinase reduces the severity of *Staphylococcus aureus* systemic infection. J Infect Dis 202:1041–9.

32. Rooijakkers SH, Ruyken M, Roos A, Daha MR, Presanis JS, Sim RB, van Wamel WJ, van Kessel KP, van Strijp JA. 2005. Immune evasion by a staphylococcal complement inhibitor that acts on C3 convertases. Nat Immunol 6:920–7.

33. Novick RP. 2003. Autoinduction and signal transduction in the regulation of staphylococcal virulence. Mol Microbiol 48:1429–49.

34. Stragier P, Kunkel B, Kroos L, Losick R. 1989. Chromosomal rearrangement generating a composite gene for a developmental transcription factor. Science 243:507–12.

35. Kunkel B, Losick R, Stragier P. 1990. The *Bacillus subtilis* gene for the development transcription factor sigma K is generated by excision of a dispensable DNA element containing a sporulation recombinase gene. Genes Dev 4:525–35.

36. Dubnau D, Losick R. 2006. Bistability in bacteria. Mol Microbiol 61:564–72.

37. Veening JW, Smits WK, Kuipers OP. 2008. Bistability, epigenetics, and bet-hedging in bacteria. Annu Rev Microbiol 62:193–210.

38. Sambrook J, Fritsch EF, Maniatis T. 1989. Molecular cloning : a laboratory manual, second edition. Cold Spring Harbor Laboratory, Cold Spring Harbor, N. Y.

39. Arnaud M, Chastanet A, Debarbouille M. 2004. New vector for efficient allelic replacement in naturally nontransformable, low-GC-content, gram-positive bacteria. Appl Environ Microbiol 70:6887–91.

40. Murphy E. 1985. Nucleotide sequence of a spectinomycin adenyltransferase AAD(9) determinant from *Staphylococcus aureus* and its relationship to AAD(3”) (9). Mol Gen Genet 200:33–39.

41. Steinmetz M, Richter R. 1994. Easy cloning of mini-Tn*10* insertions from the *Bacillus subtilis* chromosome. J Bacteriol 176:1761–3.

42. Archambaud C, Gouin E, Pizarro-Cerda J, Cossart P, Dussurget O. 2005. Translation elongation factor EF-Tu is a target for Stp, a serine-threonine phosphatase involved in virulence of *Listeria monocytogenes*. Mol Microbiol 56:383–96.

43. Livak KJ, Schmittgen TD. 2001. Analysis of relative gene expression data using real-time quantitative PCR and the 2(-Delta Delta C(T)) method. Methods 25:402–8.

44. Haque RU, Baldwin JN. 1964. Types of hemolysins produced by *Staphylococcus aureus*, as determined by the replica plating technique. J Bacteriol 88:1442–7.

45. Stokes RW, Doxsee D. 1999. The receptor-mediated uptake, survival, replication, and drug sensitivity of *Mycobacterium tuberculosis* within the macrophage-like cell line THP-1: a comparison with human monocyte-derived macrophages. Cell Immunol 197:1–9.

46. Soutourina O, Dubrac S, Poupel O, Msadek T, Martin-Verstraete I. 2010. The pleiotropic CymR regulator of *Staphylococcus aureus* plays an important role in virulence and stress response. PLoS Pathog 6:e1000894.

